# A large disordered region confers a wide spanning volume to vertebrate Suppressor of Fused as shown in a trans-species solution study

**DOI:** 10.1101/2021.06.14.447554

**Authors:** Staëlle Makamte, Amira Jabrani, Annick Paquelin, Anne Plessis, Mathieu Sanial, Aurélien Thureau, Olga Rudenko, Francesco Oteri, Marc Baaden, Valérie Biou

**Author notes:** Corresponding author: Valerie Biou.

## Abstract

Hedgehog (Hh) pathway inhibition by the conserved protein Suppressor of Fused (SuFu) is crucial to vertebrate development. By constrast, SuFu removal has little effect in drosophila.

Previous publications showed that the crystal structures of human and drosophila SuFu consist of two ordered domains that are capable of breathing motions upon ligand binding. However, the crystal structure of human SuFu does not give information about 20 N-terminal residues (IDR1) and an eighty-residue-long disordered region (IDR2) in the C-terminus, whose function is important for the pathway repression. These two IDRs are species-dependent.

We studied SuFu’s structure in solution, both with circular dichroism and small angle X-ray scattering, comparing drosophila, zebrafish and human species, to better understand this considerable difference. Our studies show that, in spite of similar crystal structures restricted to ordered domains, drosophila and vertebrate SuFu have very different structures in solution. The IDR2 of vertebrates spans a large area, thus enabling it to reach for partners and be accessible for post-translational modifications. Furthermore, we show that the IDR2 region is highly conserved within phyla but varies in length and sequence, with insects having a shorter disordered region while that of vertebrates is broad and mobile. This major variation may explain the different phenotypes observed upon SuFu removal.

## Introduction

The conserved Hedgehog (Hh) signalling pathway is essential to specify cell and tissue fate during embryogenesis and later to regulate stem cell homeostasis (Jiang and Hui, 2008). Discovered in drosophila, where it ensures proper polarisation of the larva segments (Nusslein-Volhard and Wieschaus, 1980), the Hh pathway has since been identified in many animals where it controls multiple patterning events, and has rapidly emerged as a key player in oncogenesis by controlling tumour promotion or/and progression (Pak and Segal, 2016). In the absence of Hh signal the G-Protein Coupled Receptor-related Smoothened (Smo) activity is inhibited by the Hh receptor Patched (Ptc). Binding of Hh releases this negative effect, triggering the activation of Smo, probably by favouring its association with intramembrane cholesterol (for review see (Hu and Song, 2019)). Smo activation also involves its relocalisation from intracellular vesicles to the plasma membrane in fly, and its rapid translocation from the base to the tip of the primary cilium in mammals (Hooper and Scott, 1989; Ingham et al., 1991;

Marigo et al., 1996; Zhang et al., 2018). Via its interactions with a cytoplasmic protein complex, Smo activates kinases that change the phosphorylation state of several proteins of this complex and remodel their interactions. As a result, the transcription factor(s) of the Ci/Gli family are activated and transported into the nucleus where they in turn activate the expression of specific target genes, thus changing the cell fate. The molecular backbone of the pathway is conserved but it becomes more complex from flies to mammals. For example in vertebrates, three transcription factors are involved: Gli 1, 2 and 3; their activation takes place at the primary cilium and depends on intraflagellar trafficking and results in their translocation into the nucleus (Huangfu et al., 2003).

The proper function of the Hh pathway at the right time and place is so crucial that several levels of negative regulation exist to prevent inappropriate Hh activation. This includes the products of two tumour suppressor genes: Ptc at the cell surface and Suppressor of Fused (SuFu) inside the cell. Removal of SuFu has very different consequences in drosophila and mammals. In drosophila, the phenotype of *sufu* removal is only visible in the context of the absence of *fused* kinase (Preat, 1992). Conversely, *SuFu* KO mice die in utero and exhibit heart, brain and CNS defects due to a Hh constitutional activation (Cooper et al., 2005; Svard et al., 2006).

Discovered in drosophila (Preat, 1992), where it interacts with a microtubule-associated complex including kinase Fused (Monnier et al., 1998), SuFu is a 55 kDa protein that plays an inhibitory role via the sequestration of full-length Ci in the cytoplasm. In response to Hh, this negative effect of SuFu is suppressed by the kinase Fused. SuFu then accompanies Ci into the nucleus (Sisson et al., 2006). In zebrafish, SuFu regulates Gli transfer to the primary cilium (Maurya et al., 2013) and SuFu morpholino knock down affects eye, ear and muscle development, identifying SuFu as a medium-level inhibitor of the Hh pathway (Koudijs et al., 2005). In mammals, SuFu is essential for the proper differentiation of many cell types including neural cells (Yabut et al., 2015) and it plays a role in white/brown adipocyte determination and obesity (Pospisilik et al., 2010). Here also, SuFu acts upstream of the Gli transcription factors, controlling their fate. It was thus shown to sequester full-length Gli 1 and 2 transcription factors in the cytoplasm in the absence of Hh. Moreover, SuFu also plays nuclear roles with the repressor forms of Gli as it was shown to recruit the SAP18-mSin3 repressor complex to the Gli binding sites (Paces-Fessy et al., 2004). Hh binding to Ptc represses this negative effect, allowing the Gli proteins – accompanied by SuFu-to enter the nucleus. The present state of research describes SuFu as a hub, interacting with many partner proteins, and the interactions are remodelled when Hh is activated, mostly via post-transcriptional modifications, such as phosphorylation and ubiquitylation, of SuFu (Chen et al., 2011; Raducu et al., 2016; Wang et al., 2016).

Figure S1 shows an amino acid sequence alignment of SuFu from three species belonging to different phyla: *Homo sapiens* (hSuFu), *Danio rerio* (zebrafish) (zSuFu) and *Drosophila melanogaster* (dSuFu). The crystal structures of hSuFu and dSuFu (Cherry et al., 2013; Zhang et al., 2013) harbour two ordered domains corresponding to the regions framed in blue and pink boxes in the Figure S1, with respectively 41% and 24% identity between hSuFu, zSuFu and dSuFu. The N-terminal domain is also found in prokaryotic proteins, amongst which antitoxin systems, another process involving protein-protein interactions (Zhang et al., 2011), and the C-terminal domain is specific to SuFu proteins. In addition to those two domains known from the crystal structures, Figure S1 shows that SuFu amino acid sequence contains two additional regions predicted to be intrinsically disordered by the PONDR server (http://www.pondr.com). The N-terminus disordered region IDR1 (in a dotted grey box) and the second region IDR2 (in a dotted black box) stemming from the C-terminal domains are the sequences that harbour the most striking differences between drosophila and vertebrates with 6% identical residues.

The N-terminal domain has a 7-stranded twisted beta sheet followed by a 3-helix bundle, and spans residues 30-263 (as numbered in hSuFu). The C-terminal domain has two beta sheets with 6 and 4 strands respectively and two helices; it consists of residues 267-280 and 356-457. The structure of free hSuFu (pdb code 4KM9/4BL8) is extended and the two domains become close to each other when a peptide from Gli is bound to hSuFu (pdb code 4KMD/4BLB), showing a potential tendency of the two domains to move with respect to each other. The drosophila SuFu crystal structure pdb code 4KMA spans residues 13-259 for the N-terminal domain and 265-454 for the C-terminal domain. Both domains have very similar folds to the human protein, but their relative orientations are different. The IDR2 region, which is disordered in the human structure, has only five disordered amino acids in the drosophila crystal structure. The rest of the sequence corresponding to IDR2 is structured and caps the C-terminal domain, but it has no regular secondary structure.

Due to the absence of structural information for full-length vertebrate SuFu, we used circular dichroism (CD) and small angle X-ray scattering (SAXS) to investigate the solution structures of SuFu proteins from *Drosophila melanogaster, Danio rerio* and *Homo sapiens*. We also studied hSuFuΔ30, a truncated mutant of hSuFu devoid of its IDR1 that allowed us to better analyse the effect of IDR2 alone. We found that the solution structure of the drosophila protein is very similar to the crystal structure, and investigated its breathing movements by normal mode analysis. Human and zebrafish SuFu show similar dimensions and exhibit an N-terminal IDR1 region that seems to remain close to the N-terminal domain, while the IDR2 loop in the C-terminal domain spans a large volume and can thus fetch its partner protein a long way from its core. Our present studies show that, even though their crystal structures are quite similar, the solution structures of vertebrate and drosophila SuFu are rather different and we discuss the specificity of IDR2 in the context of SuFu interaction with partner proteins.

## Experimental procedures

### 1) Protein expression and purification

dSuFu and hSuFu were cloned and expressed as previously described (Jabrani et al., 2017).

The cDNA encoding dSuFu was cloned in the pDEST17 (Invitrogen) expression vector according to standard Gateway™ protocols. The resulting plasmid encoded an N-terminal hexahistidine tag and a TEV protease cleavage site before the gene of interest. *E. coli* strain C41(DE3) (Miroux and Walker, 1996) were grown in 2YT medium at 37 °C until the optical density reached 0.6. Protein expression was induced with 0.2 mM isopropyl β-thiogalactoside overnight at 20 °C. Bacterial cells were recovered by centrifugation at 5000 g for 15 min and frozen at −80°C. Similar procedures were used for the cDNAs encoding hSuFu and zSuFu respectively, which were cloned in a pACYDuet vector. hSuFuΔ30 construct was produced by truncating the nucleotides coding for the first 30 aminoacids of hSuFu. Cells were lysed in 50 mM HEPES pH 7.5, 500 mM NaCl, 5 mM imidazole, 10% glycerol using a Constant Cell disruptor at 2.3 kbar, then centrifuged for 1 hour at 9000g. The supernatant was loaded onto a Nickel affinity column (Roche), washed in the loading buffer and eluted in a buffer containing 200mM imidazole. The eluate was collected and the buffer exchanged against 10 mM HEPES pH 7.5, 100 mM NaCl, 10 % glycerol, 2.5 mM MgCl_2_ and 2 mM β-mercaptoethanol on a concentrator. Then the protein solution was incubated with Tobacco Etch Virus protease for 24 hours at 10°C to cleave the 6-Histidine tag and the associated linker. The cleavage mixture was then dialysed back into the lysis buffer and incubated with Nickel affinity gel (Roche) for one hour to separate the cleaved protein from the TEV protease and tag. The flow through was then concentrated and injected on a Superdex 200 10/300 gel filtration column equilibrated with HEPES 50mM pH 7.5, NaCl 200mM, glycerol 10%, Dithiothreitol 5mM or bis-tris 50 mM pH 5.5, NaCl 50mM, glycerol 10% and loaded onto a HiTrapQ column, washed and eluted in a NaCl gradient. The fractions corresponding to pure SuFu protein were concentrated and used for future studies.

CD studies were performed in the protein in two different buffers: A) HEPES HCl 50mM pH 7.5, NaCl 200mM, glycerol 10%, Dithiothreitol 2mM or B) bis-tris 50 mM pH 5.5, NaCl 200mM, glycerol 10%. SAXS studies were performed in buffer A only.

### 2) SRCD measurements

Synchrotron radiation circular dichroism (SRCD) was measured at the DISCO beamline (SOLEIL synchrotron, Gif sur Yvette, France). All sample concentrations were measured using the absorbance at 280nm, after centrifugation and just prior to measurement, using a theoretical extinction coefficient of 59400 M^-1^.cm^-1^ for dSuFu, 67775 M^-1^.cm^-1^ for hSuFu and 66390 M^-1^.cm^-1^ for zSuFu. Concentrations were found to be between 4 and 10 mg/ml.

The beamline monochromator was calibrated using a camphorsulfonic acid solution prior to measurements. One microlitre of sample was deposited on one face of a calcium fluoride circular cuvette (Hellma), then the second face was carefully positioned and the cuvette closed by capillary force. Interferometry measurement showed the optical path to be 2.3 μm. The cuvette was placed in an airtight, metal sample holder that was positioned in the beamline Peltier temperature-controlled chamber, allowing for quick temperature changes with very little evaporation. Spectra were measured between 280 and 170 nm, each final spectrum being averaged from three repeated measurements. Thermal unfolding measurements were performed by averaging three spectra collected in steps of 5°C between 10 and 90 °C. Buffer spectra were taken in the same conditions for subtraction from the protein spectra. Spectra were then processed for buffer subtraction and scaling using the CDTools program (Lees et al., 2004). The melting temperature was evaluated by fitting the curve displaying the CD at 208nm with a sigmoidal function in Kaleidagraph (Synergy Software), as a function of temperature.

### 3) Small angle X-ray scattering measurements

The SAXS measurements were done at the SWING beamline (Thureau et al., submitted.) at the SOLEIL synchrotron (Gif sur Yvette, France). The sample was injected at a flux of 0.3 ml/minute into an Agilent Bio SEC-3-300 size exclusion column pre-equilibrated in the indicated buffer and mounted on an HPLC system, prior to its analysis in the X-ray beam. The X-ray energy was 12 keV. For dSuFu, measurements were carried out with an Aviex charge coupled device detector, the system being programmed to measure 100 images of the buffer then 240 images of the protein, using a 1-second exposure time interspaced by 500 ms. For hSuFu, hSuFuΔ30 and zSuFu measurements, an Eiger 4M pixel detector was used. 600 images were measured with 990ms exposure and 10 ms dead time each. The sample to detector distance is listed in Table 2.

The images were processed using the FOXTROT software (https://www.synchrotron-soleil.fr/en/beamlines/swing) developed at the SOLEIL synchrotron. The curves derived from buffer measurement were averaged and subtracted from the buffer plus protein curves. Then the radius of gyration Rg was calculated using a Guinier plot on the most intense curve and extended to all scattering curves, thus producing a plot of Rg and scattered intensity, I_0_, for all images. Curves from hSuFu and zSuFu were processed with US-SOMO (Brookes et al., 2016) to deconvolute the contribution from oligomers. The curves obtained from dSuFu did not indicate higher order species and consequently did not need deconvolution. The subtracted frames, corresponding to the deconvoluted peak with a stable Rg, were selected to produce the protein scattering curve of scattered intensity as a function of the momentum transfer q= 4πsin(Ɵ)/λ, where 2Ɵ is the diffusion angle and λ is the wavelength. Further analysis (Guinier and Kratky plots, Radius of gyration Rg and intensity at origin I_0_ calculation) was performed with the ATSAS program suite (Franke et al., 2017).

Five *ab initio* envelope calculations were carried out using the DENSS software (Grant, 2018) using the RAW interface (Hopkins et al., 2017). These were input into the DAMMAVER (Volkov and Svergun, 2003) suite to produce an averaged protein envelope.

### 4) Molecular modelling of dSuFu

The dSuFu crystal structure 4KMA was modified into an open conformation using Rosetta, (Kaufmann et al., 2010) based on the hSuFu structure (pdb ID 4KM9), as a template for relative domain orientation. This model structure, and the crystal structure 4KMA for dSuFu, were used as the initial and final frame, respectively, to guide a linear morphing obtained through the program Chimera (Pettersen et al., 2004). Each of the 20 morphing frames has been minimised without any constraint using the AMBER force field as implemented in Chimera (Pettersen et al., 2004). The 20 frames have then been analysed through the program Foxs (Schneidman-Duhovny et al., 2013) in order to calculate the corresponding scattering curve and its agreement with experimental data. The frame with the curve having the lowest chi value with respect to the experimental data was retained as the experimental model.

### 5) Molecular modelling of hSuFu missing regions

hSuFu residue numbering is according to Uniprot entry Q9VG38. Our SAXS data was measured on full-length hSuFu. The crystal structure of hSuFu PDB code 4KM9 was used as a starting point. It shows two ordered domains, but the first 20 aminoacids and 76 residues in the C-terminal domain are disordered. AllosModFoxs program (Weinkam et al., 2012) was used to build missing residues and the structure was further refined using YASARA (Krieger and Vriend, 2014) to ensure a good geometry, and the chi^2^ of the calculated scattering curve with respect to the experimental curve was 12.2. The model was then submitted to Dadimodo server https://dadimodo.synchrotron-soleil.fr (Rudenko et al., 2019) for a flexible refinement against the SAXS data by introducing flexibility in the HDR1 and the HDR2. Five models were generated and ranked according to the chi value. The two ordered domains: residues 30-255 for the N-terminal domain and residues 259-268 and 356-472 for the C-terminal domain were kept rigid while residues 1-29, 256-258 and 269-355 were left flexible. Another test was performed with a closed model starting from structure 4KMD on which the HDR1 and HDR2 regions obtained from the Yasara-refined hSuFu structure were grafted.

### 6) Molecular modelling of zSuFu

zSuFu was modelled by homology using the Phyre2 server (Kelley et al., 2015). The best-ranked model was based on hSuFu structure PDB code 4KMH. Then missing residues in the IDR2 loop (73 residues) were added with the AllosModFoxs program (Weinkam et al., 2012), yielding Chi2 values between 22 and 80. The best model was processed with Dadimodo, with residues 18 – 255 for the N-terminal and residues 259 – 268 and 356 – 472 for the C-terminal domain being kept rigid while residues 1-19, 256-258 and 279-355 were left flexible. Five models were generated and ranked according to the chi value.

## Results

### 1) Synchrotron radiation circular dichroism (SRCD) shows that truncated hSuFuΔ3O is well structured, but less stable than hSuFu

We measured SRCD spectra between 10 and 90°C for dSuFu, hSuFu, hSuFuΔ30 and zSuFu in two buffers: A) HEPES HCl 50mM pH 7.5, NaCl 200mM, glycerol 10%, Dithiothreitol 2mM or B) bis-tris 50 mM pH 5.5, NaCl 200mM, glycerol 10%. All proteins showed spectra typical of structured proteins at low temperature, and the spectra became shallower when the temperature was increased, as exemplified in Figure 1A for hSuFu. The spectra cross at an isobestic point characteristic of a two-state transition. Table 1 shows the unfolding temperatures evaluated from the variation of CD at 208 nm as a function of temperature fitted to a sigmoid function with Kaleidagraph. For hSuFu and hSuFuΔ30 the Tm was measured at pH 7.5 with DTT and at pH 5.5 without DTT. Both forms of hSuFu showed an increase in their Tm in the first condition, but hSuFu is more stabilised by pH 7.5 and DTT than hSuFuΔ30. Figure 1B shows the evaluation of secondary structure content using the BestSel program (Micsonai et al., 2015) at 10 and 60°C. At low temperature, our SuFu constructs contain between 40 and 50% of regular secondary structure and the other residues are in random coil. All studied SuFu constructs become richer in beta strand and lose helical structure when the temperature is raised from 10 to 60°C. This trend is observed in other helix-containing proteins such as myoglobin (Fändrich et al., 2003).

**Figure 1:**
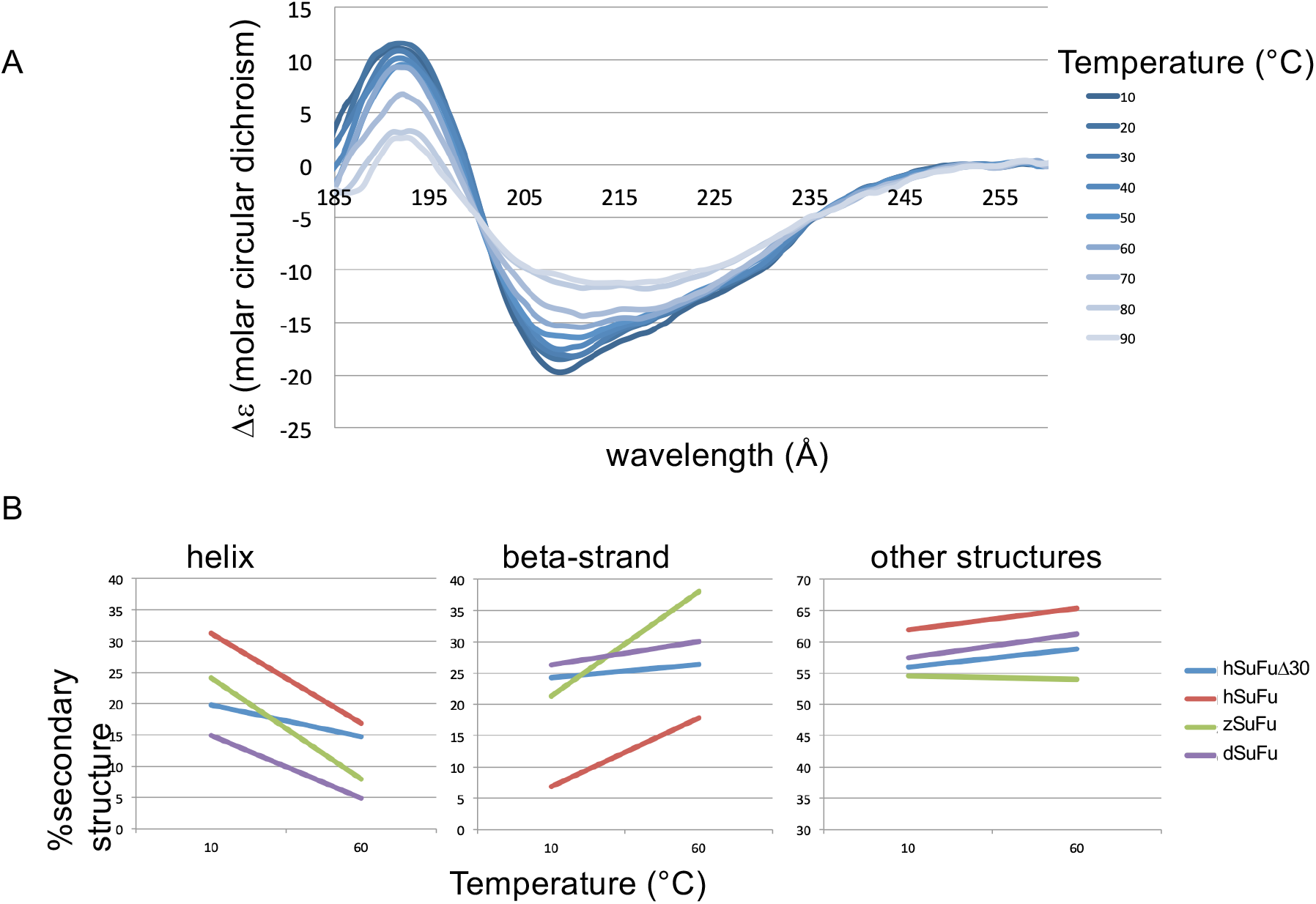
SRCD measurements of SuFu from different species. A, CD spectra showing thermal unfolding of hSuFu at pH 7.5. B, secondary structure evaluation from SRCD spectra for all studied proteins at 10 and 60°C.

**Table 1:** transition temperature for studied proteins measured by SRCD

| Protein | Buffer | T<sub>m</sub> |
| --- | --- | --- |
| zSuFu | HEPES pH 7.5, NaCl 200, glycerol 10%, DTT 2mM | 49.9° |
| hSuFu | HEPES pH 7.5, NaCl 200, glycerol 10%, DTT 2mM | 60.0° |
| hSuFu | bis-tris pH 5.5 50mM, NaCl 200, glycerol 10% | 39.1° |
| hSuFuΔ30 | HEPES pH 7.5, NaCl 200, glycerol 10%, DTT 2mM | 53.6° |
| hSuFuΔ30 | bis-tris pH 5.5 50mM, NaCl 200, glycerol 10% | 38.8° |

**Table 2:**
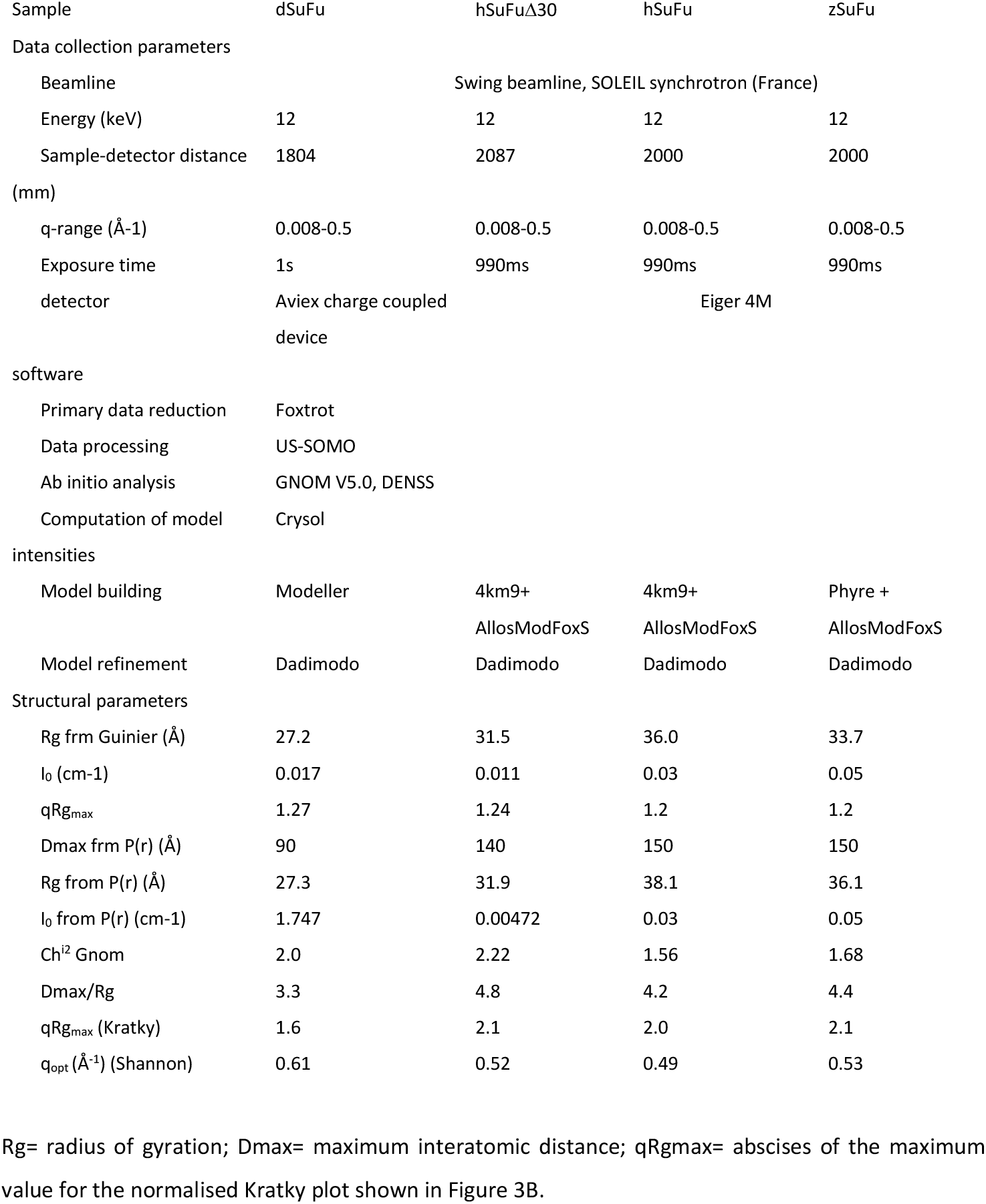
SAXS data for dSuFu, hSuFu and zSuFu. Rg= radius of gyration; Dmax= maximum interatomic distance; qRgmax= abscises of the maximum value for the normalised Kratky plot shown in Figure 3B.

### 2) SEC-SAXS and molecular modelling show that dSuFu is an elongated monomer in solution with a closed conformation

The SAXS measurements were conducted at the SWING beamline at the SOLEIL synchrotron, using a High Pressure Liquid Chromatography injection system (Thureau et al., submitted). In view of the higher stability at pH 7.5 with DTT detected with SRCD, we decided to perform our SAXS studies in those conditions. The sample migrated on the size exclusion column, and produced a peak with a homogeneous Radius of gyration (Rg) (Supplementary Figure S2) which allows us to say that the resulting SAXS curve (Figure 2A, green curve) corresponds to a monodispere particule. The estimation of the Rg using the Guinier approximation indicates an absence of aggregates and gives an Rg value of 27.2 Å. The Shannon sampling formalism indicates that the data is accurate at least up to a scattering vector of 0.4 Å^-1^. Table 2 gives a list of all measured values. The dimensionless Kratky plot (Figure 2B, grey curve) has a bell shape and does not rise at high angles, which denotes that dSuFu has a mostly compact structure with very little disorder. The distance distribution function, P(r), calculated using an indirect Fourier transform, yielded a well-behaved curve with a maximum particle dimension, D_max_, of 90Å and a maximum of the curve at r=35Å (Figure 2C, grey curve). The calculated mass of 51 kDa, obtained with Primus and Vc calculation (Rambo and Tainer, 2013), and the Rg values of 27.2 Å and 27.3 Å calculated from the Guinier and P(r) functions, respectively confirm that the particle is a monomer and is monodisperse in solution. The average envelope resulting from *ab initio* calculations encloses most of the crystal structure of dSuFu 4KMA with the exception of the 274-282 loop, which is mobile in the crystal structure, as shown by the thinner radius of the ribbon representation (Figure 3A). This figure highlights the IDR2 region that lines the C-terminal domain (residues 308-340 on the bottom left of the figure). It is ordered but it has no regular secondary structure.

**Figure 2:**
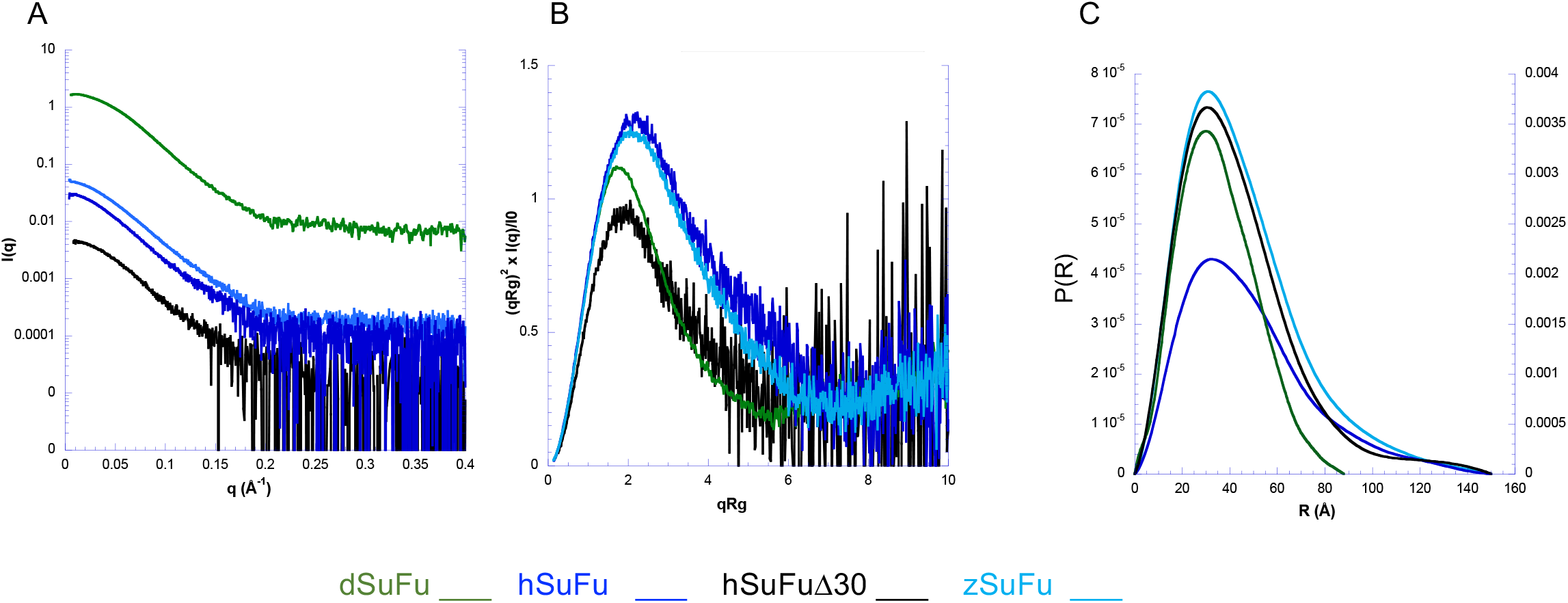
SAXS studies of hSuFu WT and Δ30, dSuFu and zSuFu. Green, dSuFu; dark blue, hSuFu WT; black, hSuFuΔ30; light blue, zSuFu. A, SAXS scattering curves B, dimensionless Kratky plots; C, pair distance distribution P(R).

**Figure 3:**
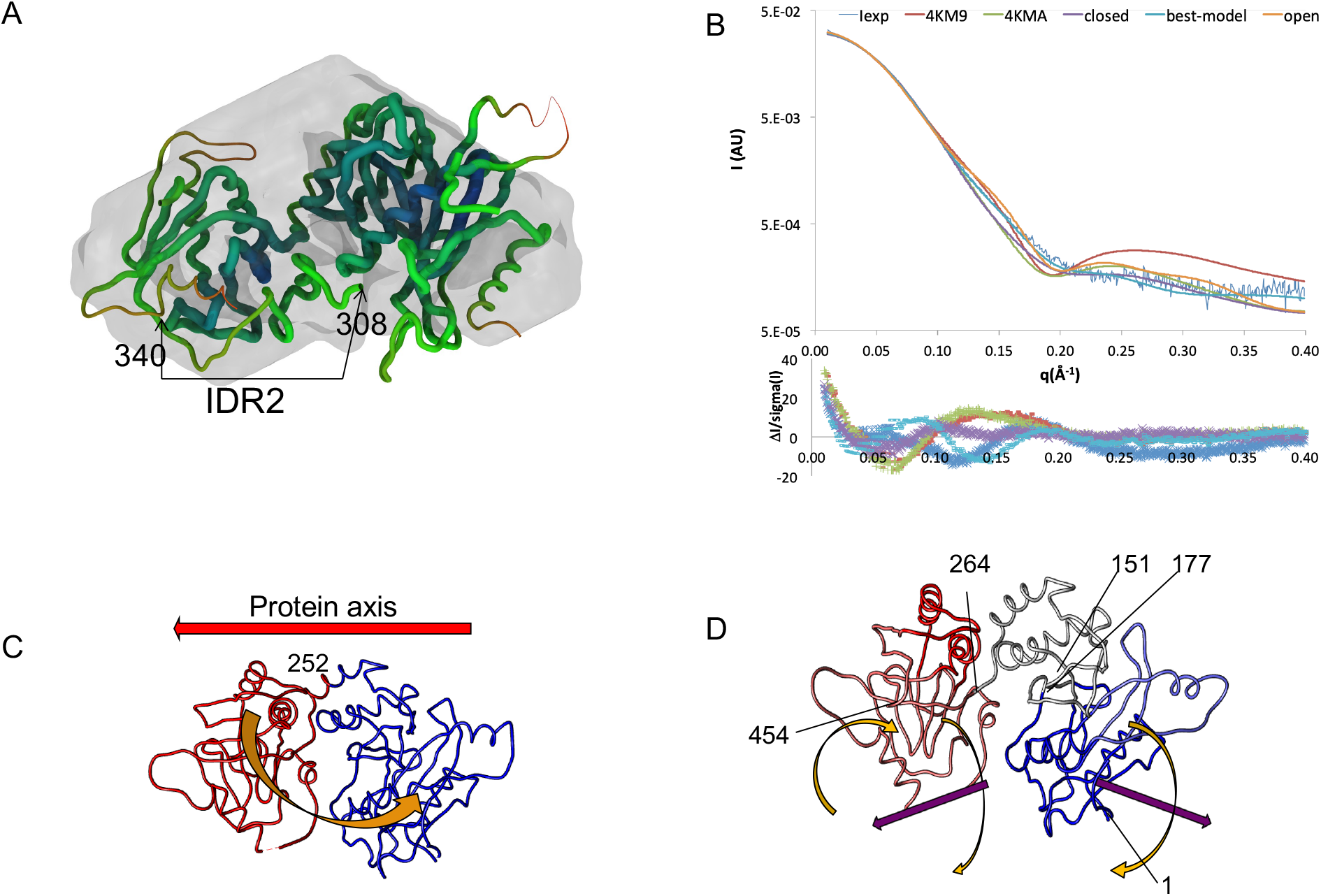
SAXS structure of drosophila SuFu. A, superposition of the crystal structure 4KMA and the envelope calculated by GASBOR and DAMAVER suite. The structure is displayed using a worm-like representation with a radius inversely proportional to the B-factor in the PDB file. B, scattering curves superposed using Crysol (top) and their differences (bottom). The experimental curve is in light blue, the PDB models 4KM9 (hSuFu) in red, 4KMA (dSuFu) in mustard green, the three models used in the normal mode simulation: closed in green, open in orange and best model in purple. C, D, the two slowest modes of movement calculated by Hingeprot. The colours distinguish the domains moving relative to each other and the arrows show the direction of movements.

The scattering curve calculated from the 4KMA crystal structure fits rather well with the SAXS spectrum, however the chi value is 7.3 and there is some disagreement between 0.15 and 0.25 Å^-1^ q value (Figure 3B, blue experimental curve, compare with 4KMA curve). We performed a modelling study in order to better characterise possible movements in dSuFu and find a model that fits the SAXS data better. The following two-step procedure was applied. First, we analysed the possible deformations in the SuFu structures. Wide rearrangements likely accounting for a conformational change in the dSuFu crystal structure have been inferred through hinge detection and compared with hSuFu crystal structures. For each structure, the Hingeprot program (Emekli et al., 2008) shows the two slowest modes of movement. For the 4KMA crystal structure, the slowest mode describes the bending of the two domains around residues Q252-D253 (Fig 3C). The second mode is described by three rigid blocks, namely N-terminal (M1 to C151 and Q177 to C194), central (Y152 to A176, and Q195 to A263) and C-terminal (G264 to E454), coloured in red, white and blue, respectively, in Figure 3D. The N- and C-terminal domains also describe a bending motion, but here the domains do not pivot on the same hinge because of the rigidity of the central block (see magenta arrows in Fig. 3D). Additionally, a wide fluctuation is observed in the loop A105-P128.

In the next step, we sampled the movements of the two domains along the direction of the bending mode in order to check if a better agreement with our SAXS results can be obtained by modelling an intermediate on this bending pathway. Two extreme open and closed conformations were generated for dSuFu, using the open (4KM9) and closed (4KMD) conformations of hSuFu as templates. The conformational change from one state to the other was modelled using linear morphing, and twenty additional intermediate conformations were generated, each of which was regularized through energy minimization, (see materials and methods section). The SAXS profiles were then simulated for each of these energy-minimised conformations and compared to the experimental profile. Chi values calculated up to a q value of 0.4 Å^-1^ are as follows: open model, 5.0; closed model, 7.7; 4KM9, 6.5; 4KMA, 7.3; best model, 3.8. The intermediate model whose diffusion curve was closest to the SAXS data was selected as the best solution (Figure 3B purple curve). This best model and the 4KMA structure of dSuFu have an RMSD of 0.7 Å for all backbone atoms. The differences mostly concern the C-terminal domain whose RMSD, with respect to the crystallized counterpart, is 0.9Å versus 0.4 Å for the N-terminal domain. Although the secondary structure is globally conserved between the proposed solution model and the crystal structure, the residues ranging from A294 – D332 in the latter are more closely packed in the domain core. The higher flexibility of these residues, containing the unresolved loop D296 – K308, accounts for a widening of the cleft between the two domains as shown by the minimum distance between the two domains (5 Å between G133-Q302 C-alpha of 4KMA structure versus 11 Å of the equivalent G147-R309 in the model). Finally, we used Dadimodo server to refine models for dSuFu starting from our best model and defined a flexible linker between the two domains consisting of residues 256-265, leaving the C-terminal domain free to move with respect to the N-terminal domain. The five resulting structures are highly similar to each other and have an average chi^2^ of 1.81±0.04 (Figure 4 AC and Table 3).

**Figure 4:**
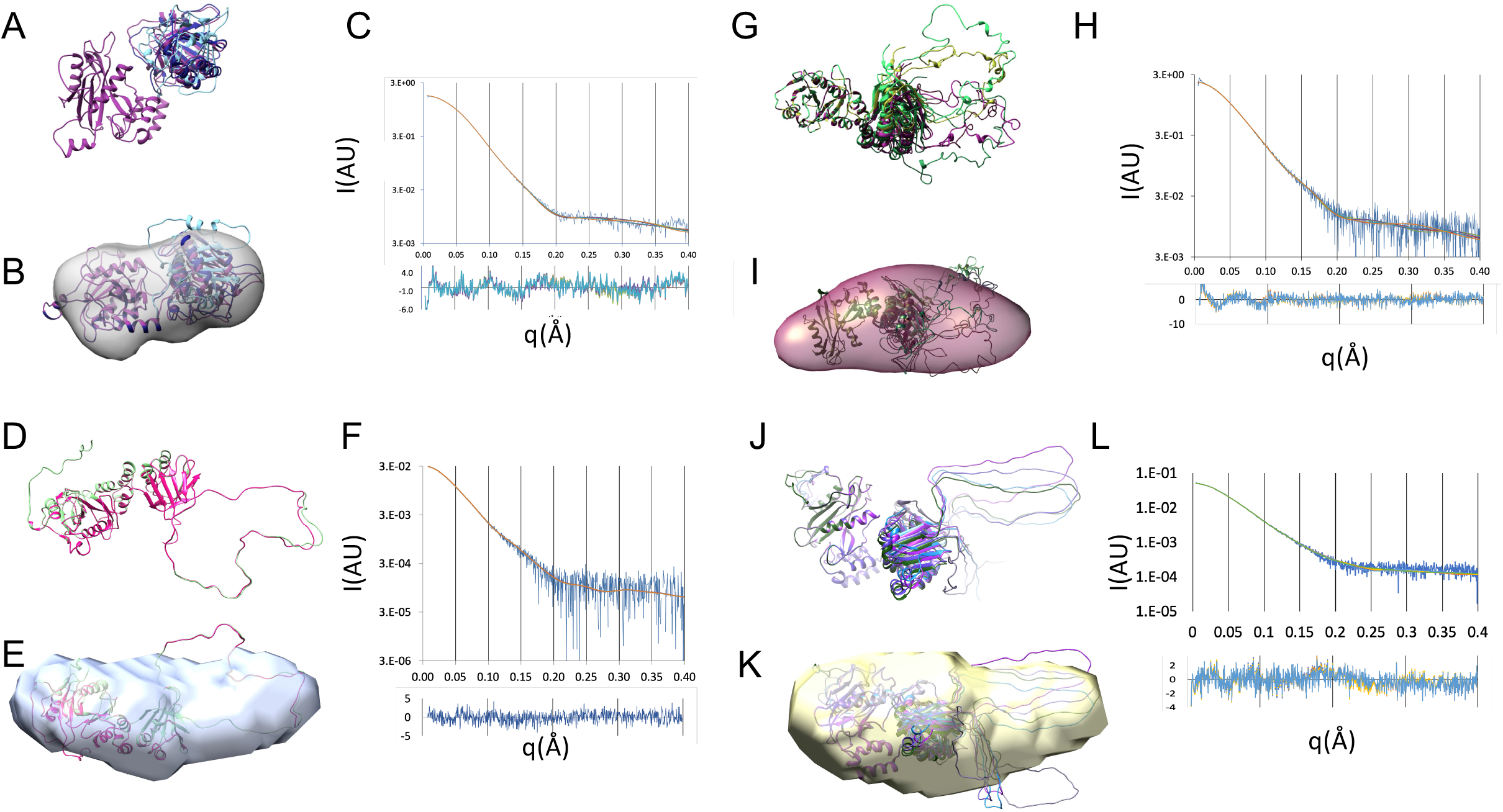
Modelling of dSuFu (A-C) hSuFu WT (D-F), hSuFuΔ30 (G-I), zSuFu (J-L) and from SAXS data. A, D, G, J: Five best models from Dadimodo; B E, I, K, rotated models superimposed with envelope from DENSS; C, F, H, L, fitting of experimental and model scattering curves.

**Table 3:** atomistic modelling of structures from SAXS data.

| sample | dSuFu | hSuFuΔ30 | hSuFu | zSuFu |
| --- | --- | --- | --- | --- |
| CRY SOL on Dadimodo output (5 structures) |  |  |  |  |
| q range ($\text{\AA}^{-1}$) | 0.008-0.4 | 0.008-0.4 | 0.008-0.4 | 0.008-0.4 |
| Chi<sup>2</sup> | 1.80±0.04 | 1.77±0.02 | 1.73±0.01 | 1.18±0.06 |
| Hydration shell contrast Dro (e- $\text{\AA}^3$) | 0.030 | 0.045 | 0.03 | 0.03 |
| Ra $\text{\AA}$ | 1.617 | 1.614 | 1.615 | 1.615 |

In summary, SAXS measurements show that the solution structure of dSuFu is slightly more open than the 4KMA dSuFu crystal structure, but is nevertheless different from the free human protein 4KM9. In addition, hinge detection shows that the clamp mode likely characterises SuFu motion. Finally, the region corresponding to IDR2 caps the C-terminal domain and is not disordered in the drosophila protein. Next, we decided to study the solution structure of vertebrate SuFu.

### 3) SEC-SAXS shows that the IDR2 disordered region spans a large volume for hSuFu and zSuFu

We chose to compare proteins from vertebrates that are far from each other in evolution: zebra fish and human. hSuFu shows 36% identity with dSuFu and 81% identity with zSuFu, while zSuFu has 35% identity with dSuFu (Fig S1). We measured SEC-SAXS data on purified hSuFu, hSuFuΔ30 and zSuFu. A reducing agent was necessary to remove a dimer-like species at pH7.5, coherent with the neutral pKa of cysteine protonation. We suspect that, at high pH, a disulphide bridge is formed involving Cysteine 292 located in the IDR2 (data not shown).

The scattering curves are shown on Figure 2A and the Radii of gyration (Rg) values are listed in Table 2. The Shannon sampling formalism shows that all curves can be used at least up to 0.4Å^-1^ (Shannon and Weaver, 1949). Associated to the dimensionless Kratky curves shown on Figure 2B, our data show that hSuFu (dark blue curve), hSuFuΔ30 (black curve) and zSuFu (tuquoise curve) span a much larger volume than dSuFu (green curve). Indeed, for comparable molecular weights, the radius of gyration is 25% higher in hSuFu and zSuFu than for dSuFu. This is in agreement with the presence of the IDR2 in zebrafish and human, which is ordered in drosophila. A comparison between the results of full-length hSuFu and truncated hSuFuΔ30 indicates the contribution of the IDR2 to the total volume. The largest interatomic distance D_max_ is 140Å, only slightly decreased by the truncation, showing that the IDR2 contributes to the majority of the D_max_ difference between drosophila (90Å) and human (150Å).

We went on to generate molecular models of SuFu whose calculated scattering curves fit the experimental curve well.

For full-length hSuFu, as explained in the Methods section, we started from free hSuFu 4KM9 and built the missing regions using AllosModFoxS (Weinkam et al., 2012). We ran YASARA (Krieger and Vriend, 2014) to regularise the geometry and then used the Dadimodo software (Rudenko et al., 2019) to find models that fit the experimental curve better. In addition, as dSuFu domains are more closed even in the absence of the Gli peptide ligand, we wanted to check if hSuFu could harbor a closed conformation in solution. We thus generated a second starting model using the Gli peptide-bound conformation of hSuFu 4KMD on which the missing regions were attached similarly to the extended model.

For the elongated models, the average chi^2^ value is 1.73 ±0.01 (Table 3). Figure 4D shows the superposition of the five models calculated. The N-terminal and C-terminal domains are in open conformation with respect to each other and the N-terminal disordered residues seem to have a rather compact structure. This is different from the IDR2 loops that protrude a long way into the solvent, conferring a very elongated structure to hSuFu. A series of *ab initio* envelopes were calculated with the DENSS program and averaged with the DAMAVER suite. The elongated envelope is in agreement with the hypothesis that IDR2 spans a large volume in solution (figures 4E). However, we consider that the most significant validation is the fit between the model and the experimental scattering curves (figure 4F). In order to further assess the conformation of the IDR2 domains we ran Dadimodo starting from the second model that had a closed conformation of the two domains. The resulting models have an average chi^2^ of 2.80±0.3. Their fit is significantly less good than those of the elongated models, showing that this set of conformations is less probable in solution. However, we cannot exclude that the protein spends part of its time in this closed conformation.

In summary, hSuFu has a very elongated conformation in solution, compatible with an open conformation of the two ordered domains and an extended IDR.

We then compared our results with those obtained with N-terminal truncated hSuFuΔ30 using the elongated YASARA model as a starting point. Dadimodo provided five models with an average chi^2^ of 1.77±0.017 (Table 3). As shown on Figure 4G, the models are slightly more variable than those of fulllength hSuFu. The C-terminal domain moves more with respect to the N-terminal one and the IDR2 loop has different relative orientations. This observation may be due to a noisier scattering curve (Figures 2A and 4H), but probably unrelated to its lower stability, as the Tm’s obtained with SRCD (see above) are comparable. Finally, The DENSS envelope has a shape similar to that of full-length hSuFu (figure 4I), in agreement with a 9% lower Dmax value.

The zebrafish SAXS data confirm that the solution structure of zSuFu is very similar to that of human SuFu, both in terms of dimensions (Table 2), models and ab initio envelope (Figure 4J-K), and significantly different from those of dSuFu. The five Dadimodo refined models have chi^2^ values of 1.18±0.06 (Figure 4L and Table 3) and have extended IDR2 loops similarly to hSuFu models.

In summary, our SAXS measurements provide structural data that the crystal structures do not supply. They show that drosophila SuFu has a very different shape in solution than vertebrate SuFu. This difference is mainly due to the presence of an extended IDR2 similar in both human and zebrafish species.

## Discussion and conclusion

Our solution investigation disclosed differences between drosophila and vertebrate SuFu that could not be anticipated from their respective crystal structures. Our SAXS results show that the closed conformation of dSuFu is not due to crystal packing effects but is intrinsic to dSuFu, and our modelling results indicate that this may be due to different hierarchy in the hinge motions, leading to a different equilibrium position. In addition, the sequence corresponding to the Intrinsically Disordered Region IDR2 is ordered in drosophila, thus stabilising the two domains in this closed conformation.

The hSuFu protein has an elongated monomeric structure and SAXS data allowed us to build models for the missing residues at the N-terminal and in the C-terminal domain. Those models suggest that the IDR2 forms an elongated loop that protrudes into the solvent. This is comforted by the study of a truncated mutant hSuFuΔ30 and of zSuFu, which is 81% identical in amino acid sequence to hSuFu and whose SAXS-derived size and shape is very similar to that of hSuFu.

The IDR2 loop thus contributes strongly to the large values of Rg and D_max_ of those proteins, and we investigated its amino acid composition in order to predict how the IDR2 of other species might behave. The IDR2 is 76 and 73 residue-long in human and zebrafish, respectively.

In the sequences aligned in Figure S1, the IDR2 starts with an aromatic residue W/F (W275 in human) belonging to the first strand of the first ß-sheet of the C-terminal domain. This aromatic establishes contacts that stabilise the interaction with conserved hydrophobic residues in the second ß-sheet. The last residues in the IDR2 are the first ones of this ß-sheet and they ensure the stabilisation of the loop ends. Disorder prediction using the Prediction of Natural Disorder PONDR server (http://www.pondr.com) shows that all sequences have a disorder tendency. Figure 5A shows the PONDR plot for the three proteins studied in this work. It confirms that dSuFu has a shorter disordered segment than zSuFu and hSuFu. In fact, if we consider the recommended disorder index threshold of 0.5, the PONDR metaserver predicts a stretch of 60 disordered residues while the crystal structure of dSuFu 4KMA shows only 11 disordered residues, confirmed by our SAXS data. Only when raising the threshold to 0.9 does the PONDR result almost predict the regions disordered in the crystal. On the other hand, with a threshold of 0.6, PONDR predicts a 90-residue-long disordered stretch for the human and zebrafish sequences, which is very close to the missing region in the crystal structures of hSuFu. This is in agreement with the observation that drosophila IDR2 region is much less disordered than the vertebrate one.

**Figure 5:**
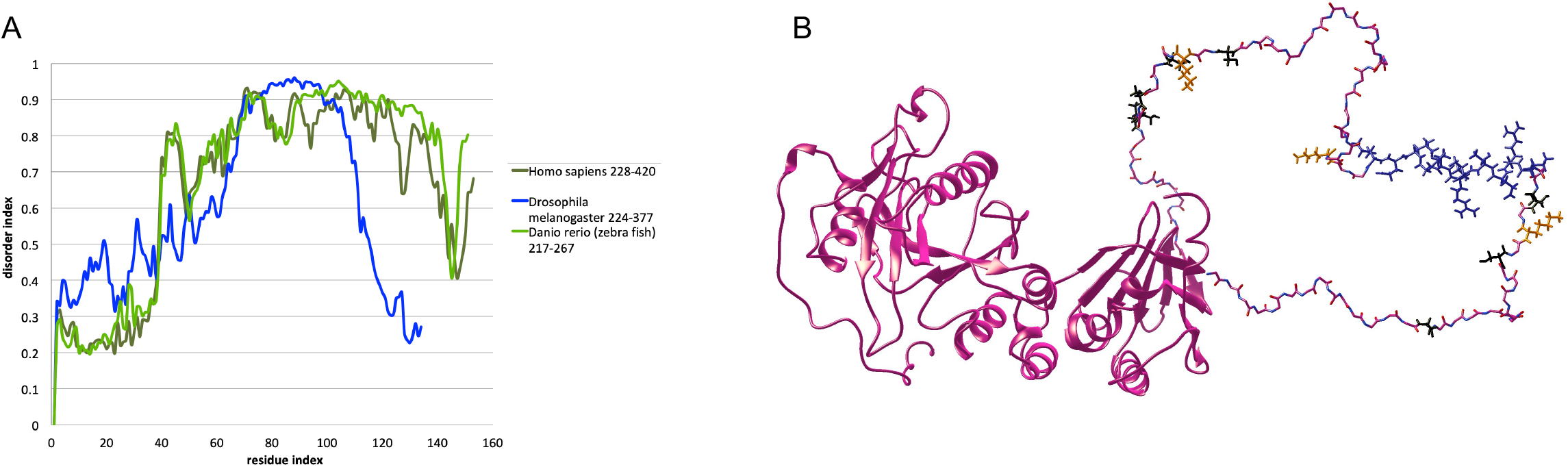
Intrinsically disordered region of SuFu: disorder prediction and structural features. A, PondR prediction of protein disorder for the three sequences studied in this work; B, hSuFu model obtained from our SAXS data showing phosphorylatable serines 290, 301, 342, 346, 349, 352 and threonines 305 and 353 are coloured black; ubiquitylated lysines 303, 321 and 344 are colored orange (Akimov et al., 2018) and nuclear export sequence (residues 308-318) is coloured blue (Zhang et al, 2017).

The elongation of IDR2 loop in vertebrates might be associated with an increased complexity in the Hh pathway from flies to vertebrates. This is exemplified by the transcription factor Cubitus interruptus (Ci) that is unique in drosophila, while three Gli proteins are simultaneously present in mammals: Gli1, Gli2 and Gli3 (Ruppert et al., 1988), allowing fine-tuning of the gene expression pattern via other partner proteins.

The function of the IDR2 region is not well known. However, Cherry et al. have studied the effect of its removal on the Hedgehog pathway activation. They found that the IDR2 is dispensable for Gli binding and Hh pathway activation with a luciferase reporter in HEK cells. However, activation of the Hh pathway in mouse embryonic fibroblast cells was repressed in the presence of full-length hSuFu but not in the presence of IDR2-truncated hSuFu. Moreover, a chimeric human protein harbouring drosophila IDR2 failed to repress the pathway (Cherry et al., 2013). This confirms that IDR2 of vertebrate SuFu has a role in the regulation of the Hh pathway, likely linked to post-translation modifications and nuclear trafficking. The amino acid composition of SuFu sequences as displayed in Table 4 shows that the IDR2, as compared to the overall sequence, is richer in acidic residues and, to a lesser extent, in basic and polar residues, and poorer in hydrophobic residues, as could be expected from the nature of IDRs. The “Phosphosite” data base (Hornbeck et al., 2004) indicates that ten out of sixteen post-transcriptional modifications of hSuFu occur in the 80 aminoacids of the IDR2. Six serine and threonine residues are predicted to be phophorylated and three lysines are predicted to be ubiquitylated or acetylated. Moreover, Zhang et al. showed that the mammalian SuFu harbours a nuclear export signal spanning residues 308-318 and two phosphorylatable serines 342 and 346 (Zhang et al., 2017). As shown on Figure 5B, those residues are located in the IRD2 making them very accessible to interact with kinases and the nuclear pore complex. This property makes the IRD2 a very good candidate to interact with exportins similarly to the complex between exportin CRM1, Ran-GTP and import adapter snurportin1 (SPN1), that involves disordered regions of SPN1 in contact with CRM1 (Monecke et al., 2009). It is known that SuFu interacts with Ci/Gli and several other partners such as SAP18, galectin3 or p66ß (Lin et al., 2014; Paces-Fessy et al., 2004). The difference between vertebrate and drosophila SuFu is further underlined by the observation that phosphorylated serines in dSuFu do not seem necessary for Hh pathway repression (Oh et al., 2015).

**Table 4:** amino acid composition of the three SuFu sequences displayed in Figure S1. The percentages are calculated using the Protparam server (https://web.expasy.org). The first column corresponds to all amino acids of the sequences; the second column corresponds to the IDR2 aminoacids as shown in the dotted black box on Figure S1. The third column is the difference between the IDR2 and the overall composition.

| aminoacid | % overall | % IDR2 | %IDR2-%overall |
| --- | --- | --- | --- |
| A | 5.38 | 5.12 | -0.26 |
| C | 1.61 | 2.33 | 0.72 |
| D | 6.50 | 8.84 | 2.34 |
| E | 7.62 | 9.30 | 1.69 |
| F | 3.84 | 1.40 | -2.45 |
| G | 7.55 | 3.26 | -4.29 |
| H | 3.14 | 1.86 | -1.28 |
| I | 4.82 | 6.51 | 1.69 |
| K | 3.91 | 6.05 | 2.13 |
| L | 10.76 | 7.44 | -3.32 |
| M | 2.10 | 1.40 | -0.70 |
| N | 3.00 | 3.72 | 0.72 |
| P | 8.04 | 8.37 | 0.34 |
| Q | 5.17 | 6.51 | 1.34 |
| R | 5.31 | 8.84 | 3.53 |
| S | 6.92 | 13.95 | 7.04 |
| T | 5.24 | 3.72 | -1.52 |
| V | 5.10 | 1.40 | -3.71 |
| W | 1.82 | 0.00 | -1.82 |
| Y | 2.17 | 0.00 | -2.17 |

In conclusion, our biophysical studies show that the disordered region of Suppressor of Fused has different lengths and properties in insects and vertebrates. In vertebrates, it is highly dynamic and spans large areas, allowing it to interact with post-translational modifying enzymes. Looking for partners specifically able to bind the IDR2 would help understand the role of this peculiar appendix.

## Acknowledgements

We are very grateful to Marie-Hélène Le Du, Dominique Durand and Patrice Vachette (Institut de Biologie Intégrative de la Cellule, CEA, CNRS, Université Paris-Saclay, France.) for help with data processing software, the SOLEIL synchrotron for access to beamlines, Javier Perez at Swing beamline, and Frank Wien and Mathieu Refrigiers at Disco beamline. Molecular graphics figures and analyses were performed with UCSF Chimera, developed by the Resource for Biocomputing, Visualization, and Informatics at the University of California, San Francisco, with support from NIH P41-GM103311.

## Author contribution

SM, AJ and APa cloned and purified proteins. SM, AJ and VB measured and processed CD and SAXS data. VB, MB and FO performed molecular modelling and analysis. AT and OR helped with Dadimodo modeling and data processing. APl and MS provided constructs and background about Hh pathway. VB designed the research, conducted research and wrote the manuscript with the help of all authors.

## Abbreviations

Ci: cubitus interruptus
Hh: Hedgehog
IDR: intrinsically disordered region
Ptc: Patched
SAXS: small angle X-ray scattering
Smo: Smoothened
SRCD: synchrotron radiation circular dichroism
SuFu: Suppressor of Fused

## Supplementary figures legends

Figure S1: alignment of SuFu aminoacid sequences from human, danio rerio and drosophila melanogaster, showing aminoacid numbering on the sides. Top line= conservation histogram; dotted grey box= IDR1; dotted black box=IDR2 for human and zebrafish; dotted red box = IDR2 for drosophila; N-terminal domain is in light blue boxes and C-terminal domain is in pink boxes. the wavy blue box shows the nuclear export signal and arrows show phosphorylable serines in the vertebrate IDR2. Green background= alpha helix; yellow background = beta strand

Figure S2: dSuFu, I_0_ and Rg SAXS profile as a function of image index.

Figure S3: hSuFu I_0_, Rg SAXS profile as a function of image index after baseline subtraction with US-SOMO.

Figure S4: zSuFu I_0_, Rg SAXS profile as a function of image index after baseline subtraction with US-SOMO.

