## Supplementary figures and images for "A large disordered region confers a wide spanning volume to vertebrate Suppressor of Fused as shown in a trans-species solution study"

### Supplementary figures

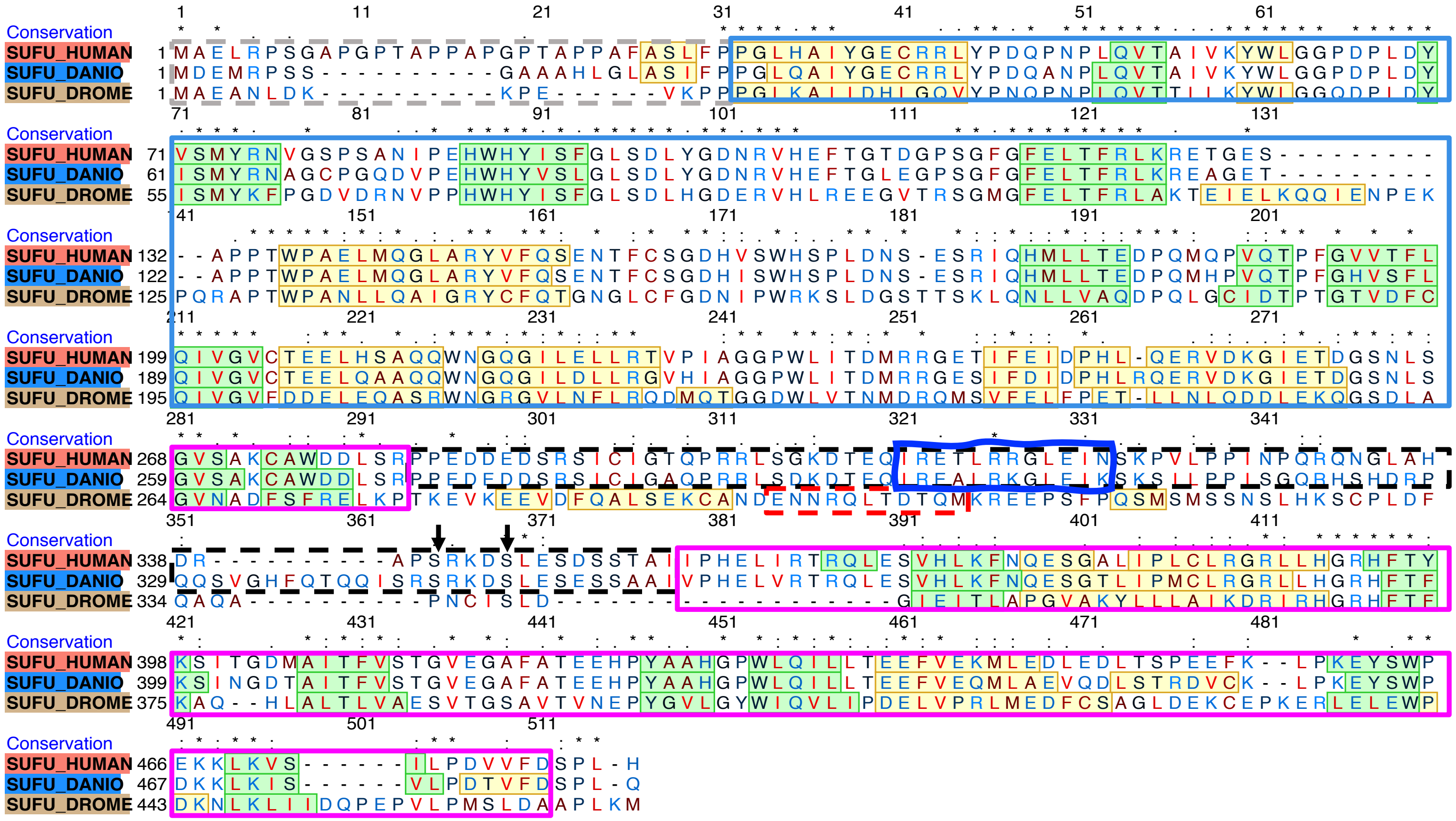

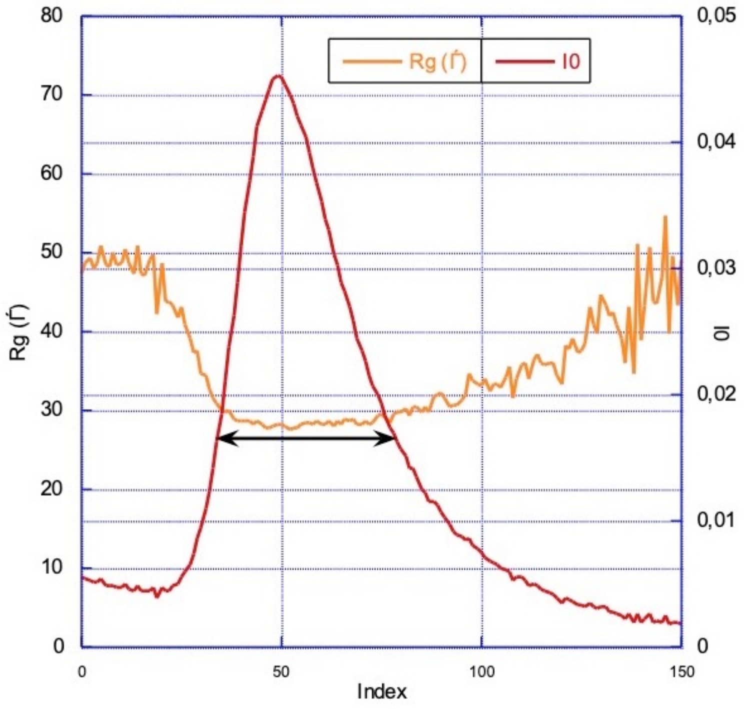

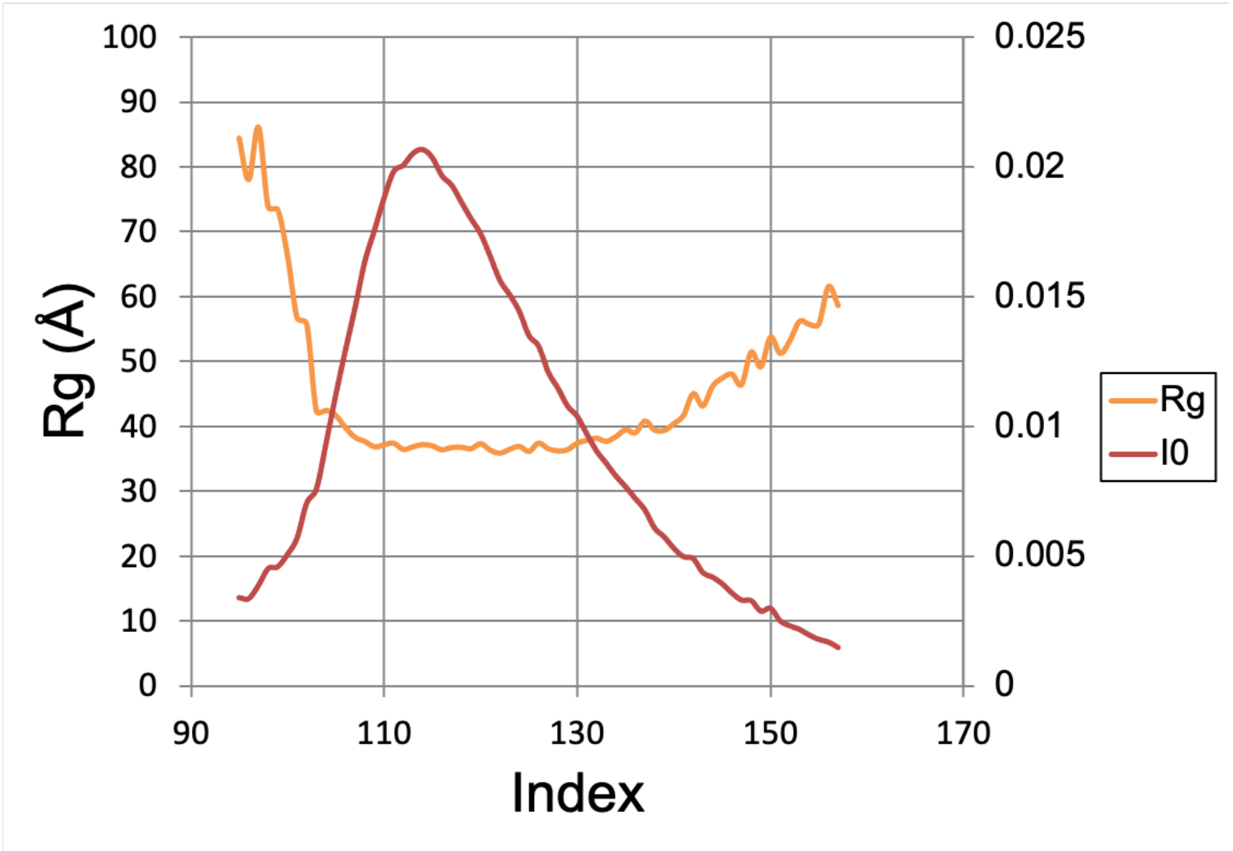

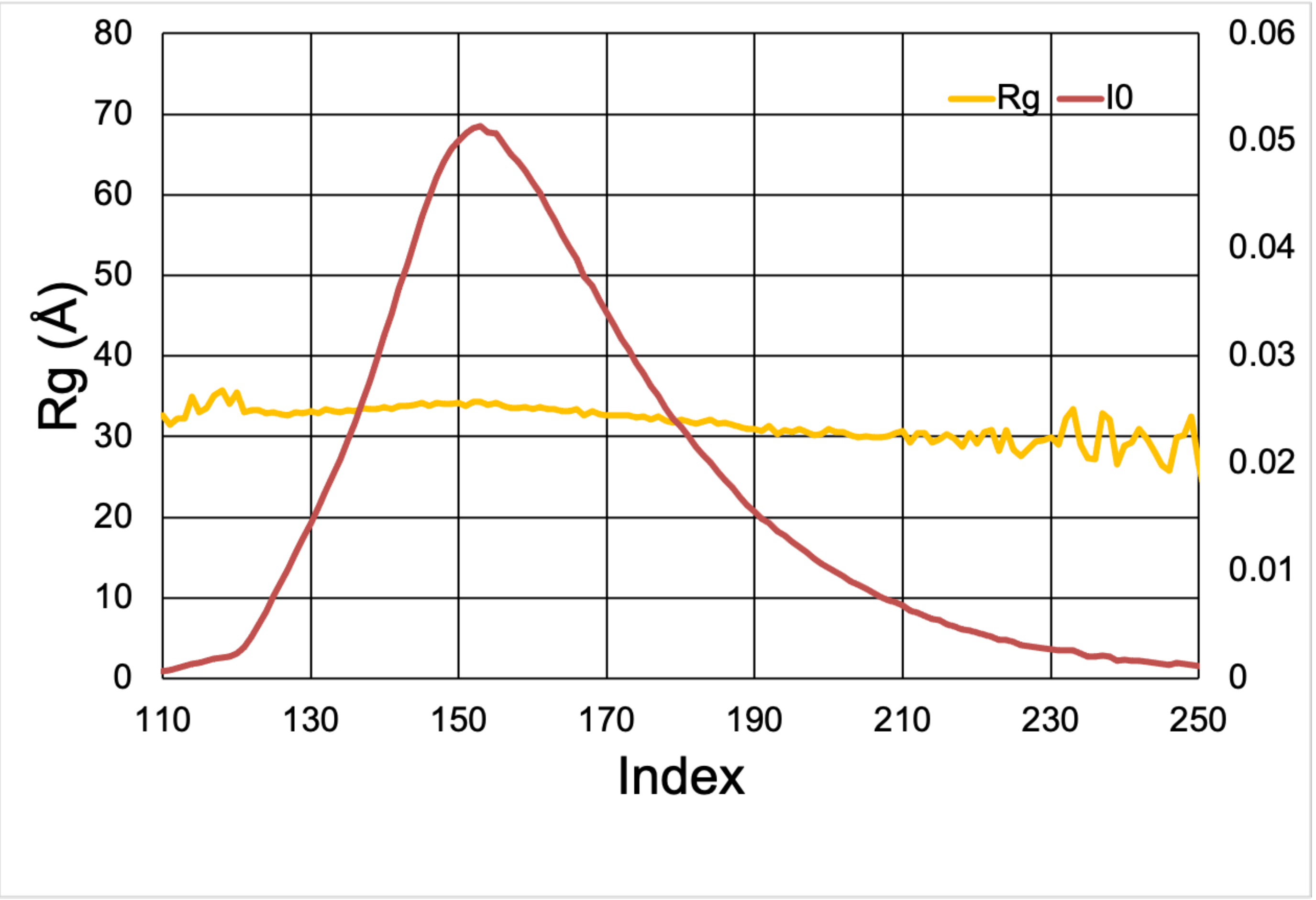
